# Diacylglycerol Kinases Balance DAG Signaling to Control B Cell Maturation, Germinal Center Homeostasis, and Humoral Immunity

**DOI:** 10.64898/2026.10.05.756823

**Authors:** Jinhai Xie, Shimeng Zhang, Wendy Lee, Huishan Tao, Yun Pan, Xunrong Luo, Xiao-Ping Zhong

## Abstract

Diacylglycerol (DAG) is a critical second messenger downstream of the B-cell receptor (BCR), but how DAG signaling is regulated during B-cell development and humoral responses remains incompletely understood. Here, complementary gain- and loss-of-function approaches revealed stage-specific requirements for diacylglycerol kinase (DGK)-regulated signaling in B cells. Expression of the gain-of-function DGKζ mutant DGKζΔNLS had limited effects on early BM B-cell development but progressively impaired transitional and mature B-cell populations. DGKζΔNLS attenuated BCR-induced ERK1/2 and mTOR signaling, activation, and proliferation and, when expressed in AID-expressing B cells, reduced GC B-cell survival and T-dependent IgG responses without detectably impairing GC B-cell proliferation in vivo. Conversely, combined DGKα/ζ deficiency resulted in marked spontaneous GC B-cell expansion associated with enhanced ERK1/2, Akt, and S6 signaling and increased glucose uptake. Despite this steady-state GC expansion, DGKα/ζ-deficient mice developed spontaneous autoreactivity and mounted impaired antigen-specific antibody responses. Thus, B-cell developmental stages exhibit differential sensitivity to altered DGK activity, and opposing perturbations of DGK-regulated signaling produce reciprocal effects on GC B-cell accumulation while both compromise productive humoral immunity. These findings identify DGKs as critical regulators that tune lipid signaling for B-cell maturation, GC homeostasis, and effective antibody responses.

## INTRODUCTION

B cells are essential mediators of humoral immunity through their ability to generate antigen-specific antibodies and immunological memory (1). B-cell development proceeds through a series of defined stages in the BM and is tightly controlled by signals from the pre-BCR and BCR (2–5). Productive rearrangement of the immunoglobulin heavy-chain locus allows expression of the μ heavy chain, which associates with the surrogate light-chain components VpreB and λ5 to form the pre-BCR and drives progression from the pro-B to pre-B stage (6–8). Subsequent light-chain rearrangement generates a functional IgM BCR, permitting progression to the immature B-cell stage. Immature B cells then migrate to the spleen, where they pass through T1 and T2 transitional stages before entering the mature FO or MZ B-cell compartments (9–11). Following antigen encounter, activated B cells can enter GCs, where they undergo clonal expansion, somatic hypermutation, affinity-dependent selection, and differentiation to generate high-affinity antibody-secreting cells and memory B cells (12–15). Thus, B cells encounter distinct signaling requirements as they progress through developmental and antigen-driven differentiation.

The pre-BCR and BCR activate multiple intracellular signaling pathways that regulate B-cell development, survival, and activation (16, 17). Following receptor engagement, Src-family kinases and Syk promote assembly of signaling complexes containing Btk, BLNK, and phospholipase Cγ2 (PLCγ2), resulting in hydrolysis of phosphatidylinositol 4,5-bisphosphate to generate the second messengers inositol-1,4,5-trisphosphate (IP3) and diacylglycerol (DAG) (18–20). Whereas IP3 initiates intracellular Ca2+ signaling, DAG recruits and activates effectors including Ras guanyl nucleotide-releasing proteins (RasGRPs) and protein kinase C (PKC) family members, thereby coupling antigen-receptor engagement to Ras–ERK, NF-κB, and other signaling pathways (21–23). Genetic disruption of PLCγ2 impairs peripheral B-cell maturation and survival, whereas combined reduction or deletion of PLCγ1 and PLCγ2 causes progressively more severe defects in early B-cell development (20, 24–26). Together with abnormalities caused by disruption of downstream DAG effectors, these findings establish an important role for PLCγ-generated second-messenger signaling during B-cell development. However, how the requirement for DAG signaling differs among developmental stages remains poorly understood.

DAG signaling is negatively regulated by diacylglycerol kinases (DGKs), a family of ten enzymes that phosphorylate DAG to generate phosphatidic acid (PA) (27). DGKα and DGKζ have important and partially overlapping functions in lymphocytes. In T cells, DGKα and DGKζ restrain TCR-induced Ras–ERK, PKC-dependent, and mTOR signaling and regulate thymocyte development, T-cell activation, regulatory T cell stability and pathogenicity, and immune responses (28–35). DGKs also regulate BCR signaling, although DGKζ appears to have a particularly prominent role in mature B cells. DGKζ deficiency markedly lowers the threshold for BCR-induced Ras–ERK1/2activation in mature FO B cells, enhances BCR-induced activation and proliferation, and increases early antibody responses to both T-independent and T-dependent antigens (23). In contrast, DGKα deficiency produced much weaker or undetectable effects on these responses, indicating that DGKζ is the predominant isoform restraining BCR-induced DAG–Ras–ERK1/2signaling under these conditions (23). Nevertheless, DGKζ deficiency causes relatively limited abnormalities in steady-state B-cell development, leaving unclear how increased DGK activity and attenuation of DAG signaling affect specific B-cell developmental transitions.

Regulation of DAG-dependent signaling may be particularly important during the GC response. GC B cells undergo repeated cycles of antigen-dependent selection, proliferation, and somatic diversification, with selected cells receiving signals that promote their survival, expansion, and differentiation (14, 36–39). BCR signaling is substantially reprogrammed in GC B cells compared with naïve B cells, and BCR-derived signals are integrated with CD40 and cytokine signals to regulate GC B-cell selection, survival, and differentiation (40–43). These features suggest that precise control of signaling may be particularly important within the GC. However, how DGK-mediated regulation of DAG signaling influences GC B-cell homeostasis and productive antibody responses remains poorly understood.

DGKζ dynamically redistributes between nuclear and cytoplasmic compartments (44). We previously generated and characterized a DGKζ mutant in which its nuclear localization sequence is disrupted (DGKζΔNLS), resulting in increased cytoplasmic retention and enhanced termination of receptor-induced DAG signaling. Expression of DGKζΔNLS strongly attenuates TCR signaling and profoundly affects thymocyte development and T-cell activation (45). Here, we used conditional expression of DGKζΔNLS to determine how enhanced DGKζ activity affects distinct stages of B-cell development and function. Vav1-iCre, CD19Cre, and AIDCre enabled analysis of DGKζΔNLS during hematopoietic development, within the B-cell lineage, and following B-cell activation, respectively. We complemented this gain-of-function approach using DGKα-deficient mice with AIDCre-mediated deletion of DGKζ to examine the consequences of reduced DGK activity in activated B cells. These complementary approaches reveal stage-specific sensitivity to altered DGK activity and reciprocal effects of enhanced DGKζ activity and combined DGKα/ζ deficiency on GC B-cell accumulation. Importantly, perturbation of DGK activity in either direction compromises productive antigen-specific antibody responses, supporting a model in which DGKs tune signaling to coordinate B-cell maturation, GC homeostasis, and humoral immunity.

## RESULTS

### Enhanced DGKζ function has limited effects on early bone marrow B cell development

DAG is a key second messenger downstream of the pre-BCR and BCR, activating signaling pathways that regulate B-cell development and activation (21, 22, 24). Previous studies have shown that DGKζ deficiency lowers the threshold for BCR-induced ERK1/2 activation in mature B cells without obviously affecting B-cell development (23). We therefore sought to determine whether increasing DGKζ activity, and thereby enhancing termination of DAG-mediated signaling, would reveal developmental requirements for DAG signaling that are not apparent following DGKζ deficiency. We previously found that DGKζΔNLS, a DGKζ mutant defective in nuclear translocation following TCR engagement, acts as a gain-of-function mutant that potently inhibits T-cell maturation and activation by attenuating TCR-induced DAG signaling(45). We therefore used an inducible Rosa26-targeted DGKζΔNLS mouse model (45) to examine the consequences of enhanced DGKζ activity during B-cell development.

To assess the effects of enhanced DGKζ function on B cell development, we crossed DGKζΔNLS mice with Vav1-iCre mice, resulting in DGKζΔNLS expression throughout the hematopoietic compartment. DGKζΔNLS is fused at its N-terminus to EGFP, allowing its expression to be monitored by GFP fluorescence (Figure 1A). Total bone marrow (BM) cell numbers were comparable between DGKζΔNLS-Vav1-iCre and control mice (WT or WT-Vav1-iCre) (Figure 1B). Within the BM, both the frequencies and absolute numbers of B220^int^CD93^+^ precursor/immature B cells and B220^hi^CD93^-^ mature B cells were similar between DGKζΔNLS-Vav1-iCre and control mice (Figure 1C, D).

**Figure 1.**
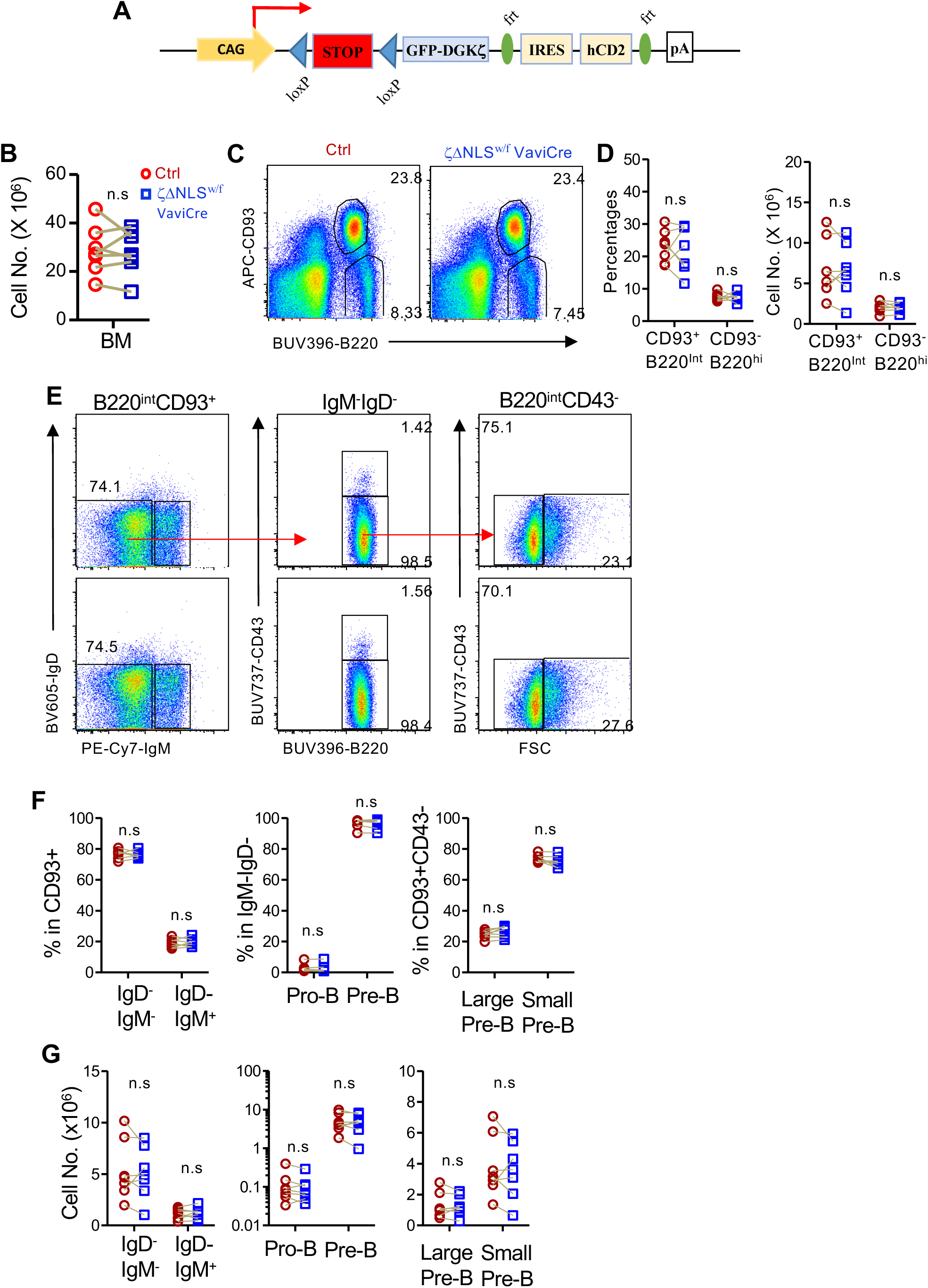
Enhanced DGKζ function has limited effects on early bone marrow B cell development. (A) Schematic illustration of the inducible DGKζΔNLS allele. (B-J) Analysis of BM B cell populations from DGKζ^ΔNLS^-Vav1^iCre^ and control mice. (B) Total BM cell numbers. (C) Representative flow cytometry plots showing B220 and CD93 expression among live BM cells. (D) Frequencies and absolute numbers of B220^int^CD93^+^ precursor/immature B cells and B220^hi^CD93^-^ mature B cells. (E) Representative flow cytometry plots and gating strategy used to identify developmental populations within B220^int^CD93^+^ BM cells. (F, G) Frequencies (F) and absolute numbers (G) of the indicated pro-B, pre-B, and immature B-cell populations. Large and small pre-B cells were distinguished on the basis of forward scatter. Data shown are representative of or pooled from seven independent experiments. Each circle or square represents one mouse, and connecting lines indicate age- and sex-matched DGKζΔNLS-Vav1-iCre and control mice analyzed in the same experiment. n.s., not significant; P < 0.05; *P < 0.01; **P < 0.001; ***P < 0.0001, determined by two tail paired Student’s t-test.

Within the B220^int^CD93^+^ compartment, pro-B cells (CD43^+^IgM^-^IgD^-^), pre-B cells (CD43^-^IgM^-^IgD^-^), including large and small pre-B cells distinguished by forward scatter, and immature B cells (IgM^+^IgD^-^) were present at comparable frequencies and absolute numbers in DGKζΔNLS-Vav1-iCre and control mice (Figure 1E-G).

Together, these findings reveal that enhanced DGKζ activity has limited effects on early BM B cell development.

### Enhanced DGKζ function impairs splenic B cell maturation and homeostasis

We next examined splenic B cell development in DGKζΔNLS-Vav1-iCre mice. Total splenocyte numbers were significantly reduced compared with control mice (Figure 2A). The frequency of B220^int^CD93^+^ immature/transitional B cells was modestly increased, whereas their absolute number was not significantly altered. The frequency and number of B220^hi^CD93^-^ mature B cells were decreased virtually in all mice except one pair of DGKζΔNLS-Vav1-iCre mice (Figure 2B, C).

**Figure 2.**
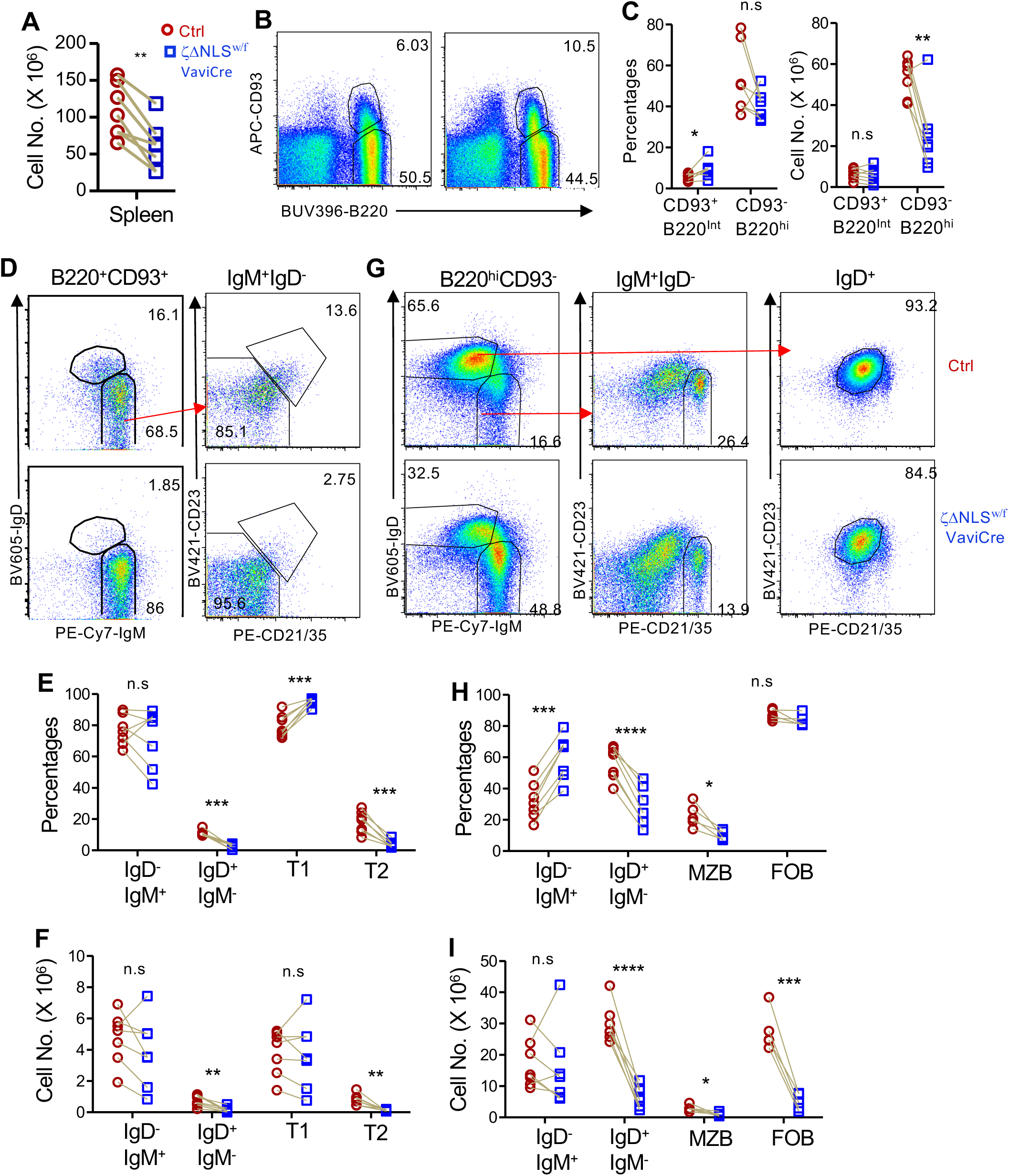
Enhanced DGKζ function impairs splenic B cell maturation. Splenocytes from DGKζΔNLS-Vav1-iCre and control mice were analyzed by flow cytometry. (A) Total splenocyte numbers. (B) Representative flow cytometry plots showing B220 and CD93 expression among live splenocytes. (C) Frequencies and absolute numbers of B220^int^CD93^+^ immature/transitional B cells and B220^hi^CD93^-^ mature B cells. (D) Representative flow cytometry plots and gating strategy for analysis of B220^int^CD93^+^ transitional B cells. IgM^+^IgD^-^ transitional B cells were further analyzed for CD21 and CD23 expression to identify CD21^-^ CD23^-^ T1 and CD21^+^CD23^+^ T2 B cells. (E, F) Frequencies (E) and absolute numbers (F) of the indicated IgM/IgD-defined populations and T1 and T2 B cells. (G) Representative flow cytometry plots and gating strategy for analysis of B220^hi^CD93^-^ mature B cells. IgM^+^IgD^-^ and IgD^+^ populations were identified, and mature B cells were further analyzed for CD21 and CD23 expression to identify MZ and FO B cells. (H, I) Frequencies (H) and absolute numbers (I) of the indicated mature B-cell populations. Data shown are representative of or pooled from seven independent experiments. Each circle or square represents one mouse, and connecting lines indicate age- and sex-matched DGKζΔNLS-Vav1-iCre and control mice analyzed in the same experiment. n.s., not significant; P < 0.05; *P < 0.01; **P < 0.001; ***P < 0.0001, determined by paired Student’s t-test.

Within the B220^int^CD93^+^ transitional B-cell compartment, the frequency and number of IgM^+^IgD^-^ transitional B cells were not significantly altered in DGKζΔNLS-Vav1-iCre mice, whereas IgD^+^ cells were substantially reduced (Figure 2D-F). Transitional B cells were further resolved into CD21^-^CD23^-^ T1 and CD21^+^CD23^+^ T2 cells. DGKζΔNLS-Vav1-iCre mice displayed an increased proportion of T1 cells, whereas their absolute numbers remained comparable to controls. In contrast, T2 cells were significantly reduced in both frequency and absolute number (Figure 2D-F). Thus, enhanced DGKζ activity appeared to preferentially impair progression from the T1 to T2 stage rather than the generation or accumulation of T1 B cells.

The marked reduction in the absolute number of splenic B220^hi^CD93^-^ mature B cells in DGKζΔNLS-Vav1-iCre mice further suggested that enhanced DGKζ activity exerted a stronger effect on late B cell maturation and/or homeostasis than on early B cell development. Within the B220^hi^CD93^-^ compartment, the relative frequency of IgM^+^IgD^-^ cells was increased in DGKζΔNLS-Vav1-iCre mice, although their absolute number was not significantly altered. In contrast, IgM^-^IgD^+^ cells were markedly decreased in both frequency and absolute number (Figure 2G-I). Within splenic mature IgM^+^IgD^-^ cells, CD21^+^CD23^-^ marginal zone (MZ) cells were decreased in both frequency and absolute number (Figure 2G-I) and the number of CD21^-^ CD23^+^ follicular (FO) B cells within the IgD^+^ mature B-cell population was markedly reduced.

Together, these results demonstrate that enhanced DGKζ activity has relatively little effect on the generation of early B-cell precursors and T1 B cells but impairs subsequent splenic B-cell maturation and/or maintenance. The pronounced reductions in T2, MZ, and FO B-cell populations identify transitional and mature stages of B-cell development as particularly sensitive to attenuation of DAG-mediated signaling.

### Enhanced DGKζ activity intrinsically impairs B cell maturation

Because DGKζΔNLS is expressed throughout the hematopoietic compartment in DGKζΔNLS-Vav1^iCre^ mice, the impaired B cell maturation observed in these mice could result either from effects within B cells or indirectly from alterations in other hematopoietic cells. To determine whether these developmental defects were cell intrinsic, we generated mixed BM chimeras by reconstituting lethally irradiated CD45.1^+^ WT recipient mice with a 1:1 mixture of BM cells from CD45.2^+^ WT and DGKζΔNLS-Vav1iCre donors. Because DGKζΔNLS is fused to EGFP, WT and DGKζΔNLS-expressing donor-derived cells could be distinguished as GFP- and GFP^+^ cells, respectively, within the CD45.2^+^CD45.1^-^ compartment (Figure 3A). This competitive chimera approach allowed WT and DGKζΔNLS-expressing B cells to develop within the same hematopoietic environment.

**Figure 3.**
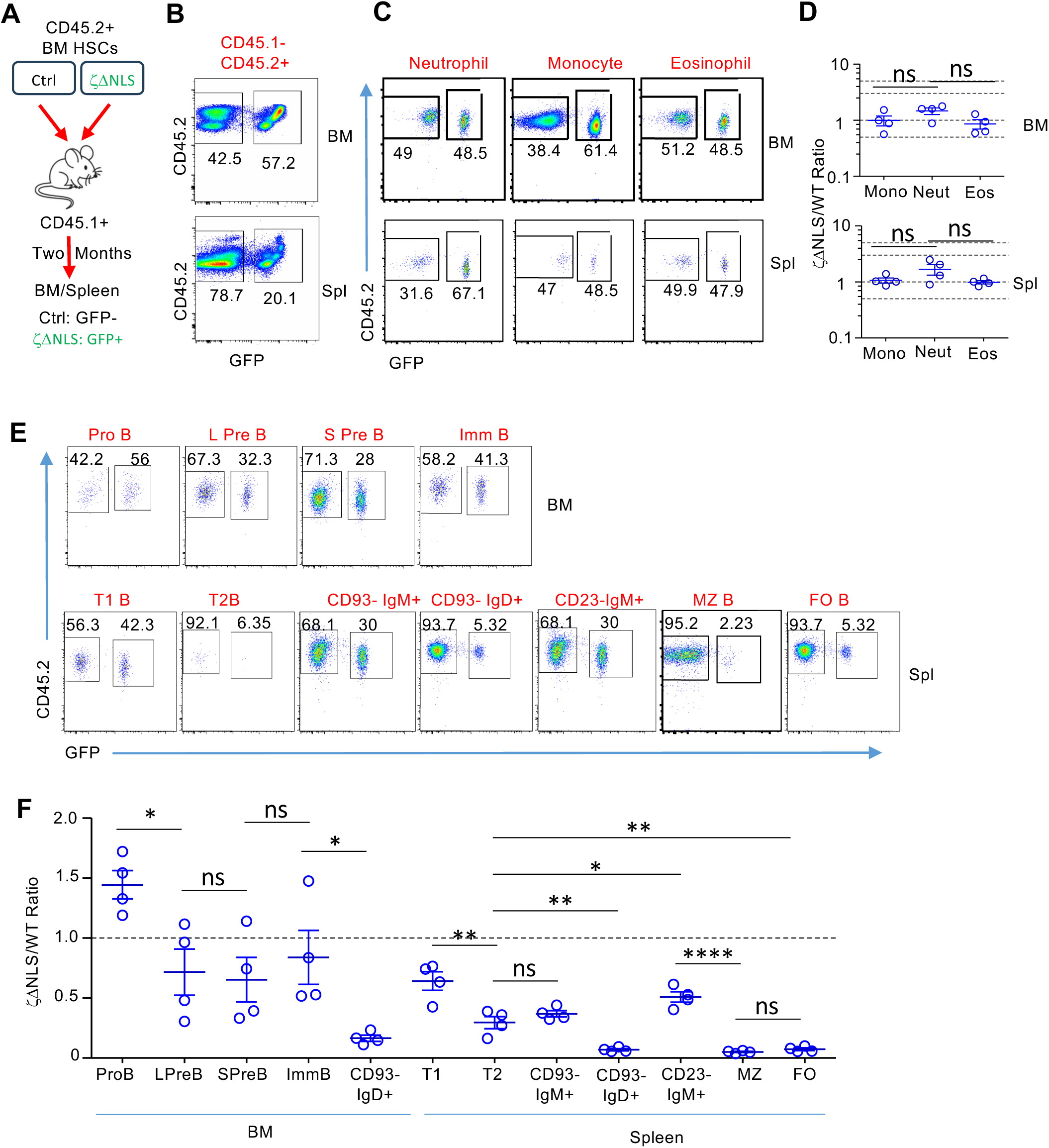
Intrinsic inhibition of B cell maturation by DGKζΔNLS. (A) Experimental design. Lethally irradiated CD45.1^+^ WT recipient mice were reconstituted with a 1:1 mixture of BM cells from CD45.2^+^ WT (GFP^-^) and DGKζΔNLS-Vav1^iCre^ (GFP^+^) donor mice. BM and spleen cells were analyzed two months after reconstitution. (B) Identification of WT (GFP^-^) and DGKζΔNLS-expressing (GFP^+^) cells among live CD45.2^+^CD45.1^-^ donor-derived BM cells and splenocytes. (C) Representative FACS plots showing percentages of GFP^+^ and GFP^-^ populations within monocytes, neutrophils, and eosinophils. (D) Scatter plot show Mean ± SEM of DGKζΔNLS/WT (GFP^+^/GFP^-^) ratios of BM and splenic monocytes, neutrophils, and eosinophils in individual recipient mice. (E) Representative FACS plots showing percentages of GFP^+^ and GFP^-^ populations during B cell development and maturation. (F) Scatter plot show Mean ± SEM of DGKζΔNLS/WT (GFP^+^/GFP^-^) ratios of indicated B cell populations in individual recipient mice. Each symbol represents one recipient mouse. Data shown represent or are pooled from three independent experiments.

Two months after reconstitution, CD45.1^-^CD45.2^+^BM cells and splenocytes from the recipients could be separated into DGKζΔNLS-derived (GFP^+^) to WT-derived (GFP^-^) populations (Figure 3B). The GFP^+^/GFP^-^ ratios of monocytes (CD11b^+^Ly6C^+^) and eosinophils (CD11b^+^Siglec-F^+^) were close to 1 in both the BM and spleen (Figure 3C,D), indicating comparable contributions of the two donor populations to these myeloid lineages. Although the average GFP^+^/GFP^-^ ratios of neutrophils was close to 2, it was not statistically different from monocytes and eosinophils.

In the BM, this ratio was approximately 1.4 in pro-B cells but decreased to approximately 0.7 in both large and small pre-B cells (Figure 3E,F). This reduction suggests that enhanced DGKζ activity conferred a modest competitive disadvantage during the pro-B to pre-B transition, an effect that was not readily apparent under the noncompetitive conditions in DGKζΔNLS-Vav1^iCre^ mice. This ratio was slightly increased to 0.8 in IgM^+^ immature B cells but drastically reduced to 0.16 in CD93^-^IgD^+^ mature B cells. These data further suppor that enhanced DGKξ activity exerts weak or minimal effects on early B cell development in the BM.

In the spleen, the DGKζΔNLS/WT ratio of T1 cells was close 0.64, which was not significantly different from that of immature B cells in the BM. However, the ratio in T2 cells was reduced to 0.3 and was statistically lower than that of T1 (p<0.01), consistent with impaired progression through the T1-to-T2 transition. The competitive disadvantage was also noted in overall IgM^+^ mature B cells and became substantially greater among MZ and FO B cell populations. DGKζΔNLS-expressing MZ and FO B cells were markedly depleted relative to their WT counterparts (Figure 3E,F). These findings indicate that the generation and/or maintenance of mature MZ and/or FO B cells is particularly sensitive to enhanced DGKζ activity.

Together, these competitive chimera experiments demonstrate that the effects of enhanced DGKζ activity on B cell maturation are cell intrinsic and vary substantially across developmental stages. DGKζΔNLS caused a modest competitive disadvantage during the pro-B to pre-B transition but had relatively limited effects on subsequent early BM development. In contrast, the disadvantage became progressively more pronounced during peripheral maturation, beginning with the T1-to-T2 transition and culminating in the marked depletion of MZ and FO B cells. These findings indicate that distinct stages of B cell development differ in their sensitivity to enhanced DGKζ activity and suggest that mature peripheral B-cell populations are particularly dependent on appropriately sustained DAG signaling.

### Enhanced DGKζ activity attenuates BCR signaling and inhibits BCR- and LPS-induced B cell proliferation

Having established that enhanced DGKζ activity intrinsically impairs B cell maturation, we next sought to determine how DGKζΔNLS affects BCR-induced signaling and functional responses. For these experiments, we crossed DGKζΔNLS mice with CD19^Cre^ mice (46) to induce DGKζΔNLS selectively within the B cell lineage. In contrast to the broadly expressed Vav1^iCre^ model used to assess B cell development, this approach allowed us to examine the effects of DGKζΔNLS more directly within B cells. Because CD19^Cre^-mediated recombination was incomplete, DGKζΔNLS-CD19^Cre^ mice contained both GFP^+^ DGKζΔNLS-expressing and GFP^-^ non-recombined B cells, providing an internal control for assessing the effects of DGKζΔNLS expression (Figure 4B).

**Figure 4.**
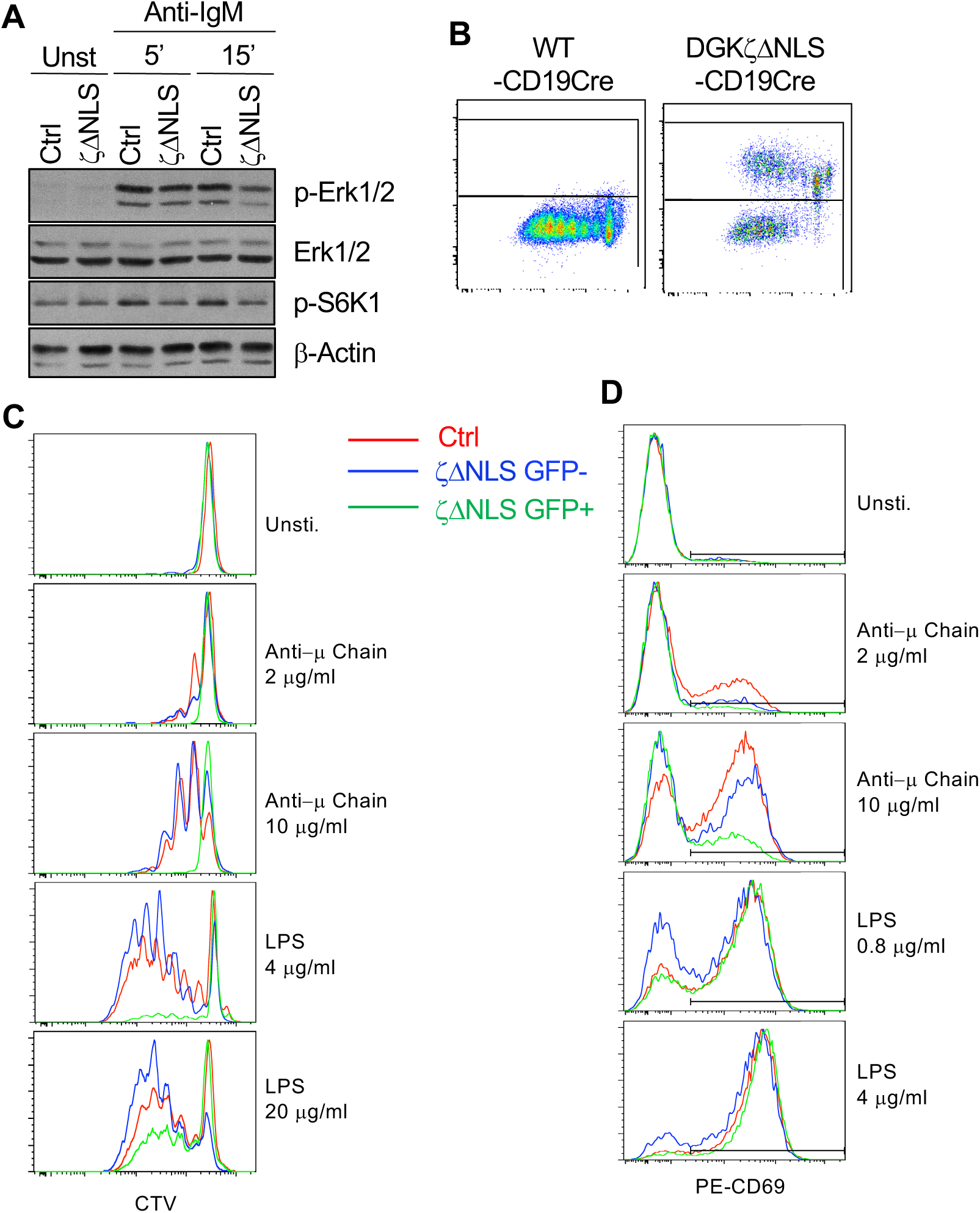
Enhanced DGKζ activity attenuates BCR signaling and inhibits BCR- and LPS-induced B cell proliferation. Splenic B cells from DGKζΔNLS-CD19^Cre/+^ and WT-CD19^Cre/+^ (Ctrl) mice were analyzed for BCR- and LPS-induced signaling, proliferation, and activation. (A) Analysis of BCR-induced signaling. Splenic B cells were left unstimulated (Unst) or stimulated with anti-IgM for 5 or 15 min. Cell lysates were analyzed by immunoblotting for phosphorylated ERK1/2 (p-ERK1/2), total ERK1/2, phosphorylated S6K1 (p-S6K1), and β-actin. (B, C) Analysis of B cell proliferation. CTV-labeled splenocytes were left unstimulated or stimulated for 80 h with the indicated concentrations of anti-IgM or LPS. (B) Representative flow cytometric plots showing the identification of GFP^-^ and GFP^+^ B cells from DGKζΔNLS-CD19^Cre^ mice. (C) Representative overlaid histograms showing CTV dilution in control B cells (red), GFP^-^ B cells from DGKζΔNLS-CD19^Cre^ mice (blue), and GFP^+^ DGKζΔNLS-expressing B cells from the same mice (green). (D) Analysis of B cell activation. Representative overlaid histograms showing CD69 expression following overnight stimulation with the indicated concentrations of anti-IgM or LPS. Colors denote the same B-cell populations as in (C). Data are representative of three independent experiments.

We first examined BCR-induced signaling following anti-IgM stimulation. Anti-IgM induced robust ERK1/2 and S6K1 phosphorylation in control B cells, whereas phosphorylation of both proteins was attenuated in B cells from DGKζΔNLS-CD19^Cre^ mice (Figure 4A). These findings are consistent with important roles of DAG signaling in mTOR signaling and enhanced DGKζ activity suppressing DAG-dependent signaling to both the Ras–ERK1/2and mTOR pathways, as we previously observed in T cells (31, 45, 47). Because immunoblotting was performed using total B cells containing both GFP^+^ recombined and GFP^-^ non-recombined cells, incomplete induction of DGKζΔNLS likely resulted in an underestimation of its inhibitory effects on BCR signaling.

We next determined whether enhanced DGKζ activity affected B cell proliferation. CTV-labeled splenocytes from control and DGKζΔNLS-CD19-Cre mice were stimulated for 80 h with anti-IgM or LPS, and proliferation was assessed by CTV dilution after separating GFP^+^ DGKζΔNLS-expressing and GFP^-^ non-recombined B cells (Figure 4B). Following anti-IgM stimulation, GFP^+^ DGKζΔNLS-expressing B cells showed markedly reduced proliferation compared with both control B cells and GFP^-^ B cells from the same cultures, with the inhibitory effect particularly evident at the higher concentration of anti-IgM. Unexpectedly, DGKζΔNLS also inhibited LPS-induced proliferation. This inhibition was most apparent at the lower LPS concentration and was partially overcome by increasing the concentration of LPS. Thus, enhanced DGKζ activity inhibits proliferation induced not only through the BCR but also through TLR4.

To determine whether DGKζΔNLS also affected an earlier B cell activation response, we measured surface CD69 expression following overnight stimulation (Figure 4C). Anti-IgM-induced CD69 upregulation was markedly attenuated in GFP^+^ DGKζΔNLS-expressing B cells compared with control and GFP^-^ B cells. In contrast, LPS induced robust CD69 upregulation in GFP^+^ DGKζΔNLS-expressing cells, with little difference from control or GFP^-^ cells. Thus, although enhanced DGKζ activity impaired LPS-driven proliferation, it did not broadly suppress early activation induced through TLR4.

Together, these results demonstrate that enhanced DGKζ activity attenuates BCR-induced ERK1/2signaling and strongly inhibits BCR-driven activation and proliferation. DGKζΔNLS also inhibited LPS-induced proliferation despite largely preserved LPS-induced CD69 upregulation, indicating that its effects are not restricted to BCR signaling. These findings suggest that DGK-regulated lipid signaling contributes to proliferative responses downstream of TLR4, while being less critical for at least some early TLR4-induced activation events.

### Enhanced DGKζ activity impairs GC B cell survival and antigen-specific IgG responses

The strong inhibition of BCR-induced activation by DGKζΔNLS prompted us to investigate whether enhanced DGKζ activity also affects B cell responses following antigen activation. GC B cells undergo iterative rounds of somatic diversification, selection, and proliferation that drive affinity maturation and ultimately generate high-affinity antibody-secreting cells and memory B cells (14). BCR signaling is dynamically regulated and substantially rewired during the GC response (40, 41), raising the possibility that tight control of DAG-mediated signaling may be particularly important for GC B cell responses. However, analysis of GC responses using Vav1-iCre or CD19-Cre would be complicated by the effects of DGKζΔNLS on earlier B cell development and mature B cell populations. We therefore used AID-Cre (48) to induce DGKζ constructs in activated B cells, allowing us to assess the effects of enhanced DGKζ activity during the GC response while minimizing effects on earlier stages of B cell development.

We first examined B cell populations in EGFP-DGKζΔNLS-AID^Cre/+^ and WT-AID^Cre/+^ mice under steady-state conditions. Consistent with the later onset of AID-Cre activity, total cell numbers in the BM, spleen, and Peyer’s patches (PPs), as well as the frequencies and numbers of total B220^+^ B cells and major IgM/IgD-defined B cell populations, were not obviously altered in EGFP-DGKζΔNLS-AID^Cre/+^ mice compared with controls (Supplemental Figure S1). The frequencies and numbers of GL7^+^Fas^+^ GC B cells within the B220^+^IgM^-^IgD^-^ population in the spleen and PPs were also not significantly different between EGFP-DGKζΔNLS-AID^Cre/+^ and control mice, although they showed a trend toward reduction in EGFP-DGKζΔNLS-AID^Cre/+^ mice (Figure 5A).

**Figure 5.**
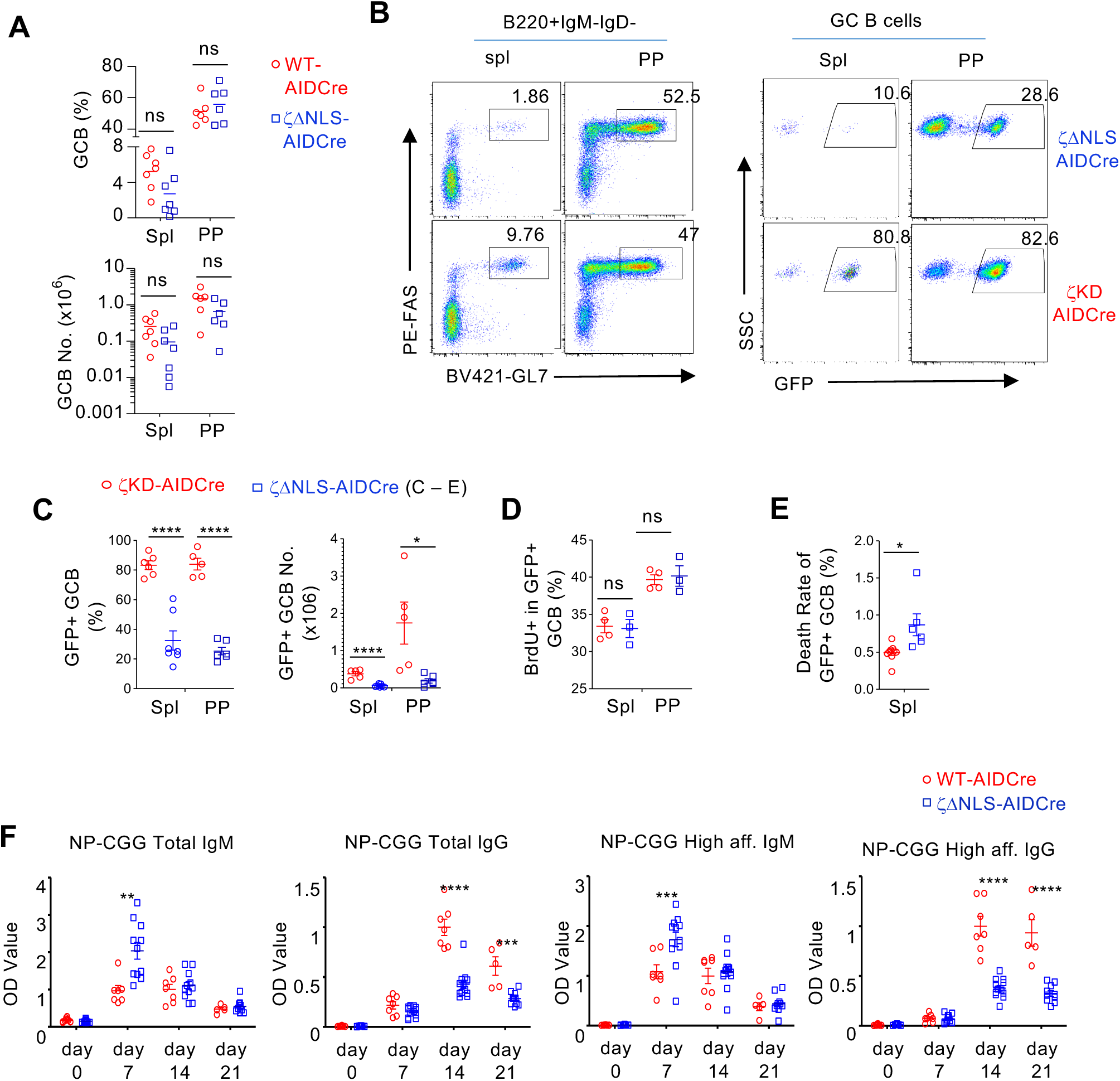
Enhanced DGKζ activity in activated B cells impairs GC B cell survival and antigen-specific IgG responses. (A) Frequencies and absolute numbers of GL7^+^Fas^+^ GC B cells in the spleen and Peyer’s patches (PPs) of 8- to 10-week-old EGFP-DGKζΔNLS-AID^Cre/+^ and WT-AID^Cre/+^ mice. (B–E) Splenocytes and PP cells from EGFP-DGKζΔNLS-AID^Cre/+^ and EGFP-DGKζKD-AID^Cre/+^ mice were analyzed. (B) Representative flow cytometric plots showing GL7 and Fas expression among live B220^+^IgM^-^IgD^-^ B cells and GFP expression within GC B cells. (C) Frequencies and absolute numbers of GFP^+^ cells within the GC B cell population. (D) Frequencies of BrdU^+^ cells among GFP^+^ GC B cells 16 h after intraperitoneal injection of BrdU. (E) Death rates of GFP^+^ GC B cells. (F) Serum total and high-affinity NP-specific IgM and IgG titers in EGFP-DGKζΔNLS-AID^Cre/+^ and WT-AID^Cre/+^ mice at 7, 14, and 21 days after intraperitoneal immunization with NP-CGG in alum. Total and high-affinity NP-specific antibodies were measured using NP4-BSA and NP26-BSA, respectively. Each symbol represents one mouse. Data are representative of or pooled from at least three independent experiments and are presented as mean ± SEM where indicated. n.s., not significant; P < 0.05; P < 0.01; P < 0.001; P < 0.0001, as determined by unpaired two-tailed Student’s t-test.

Because AID-Cre-mediated recombination induces expression of EGFP-DGKζΔNLS, GFP expression allowed us to identify GC B cells in which the DGKζΔNLS transgene had been induced. Notably, GFP^+^ cells were poorly represented within the GC B cell population of EGFP-DGKζΔNLS-AID^Cre/+^ mice (Figure 5B, C). To determine whether this depletion depended on enhanced DGKζ catalytic activity, we compared these mice with EGFP-DGKζKD-AID^Cre/+^ mice, which inducibly express an EGFP-fused kinase-dead DGKζ mutant(45). In contrast to DGKζΔNLS-expressing cells, GFP^+^ DGKζKD-expressing cells were readily detected within the GC B cell compartment (Figure 5B, C). Thus, the selective depletion of GFP^+^ DGKζΔNLS-expressing GC B cells depended on DGKζ kinase activity rather than simply expression of the DGKζ transgene or EGFP.

We next asked whether the reduced representation of DGKζΔNLS-expressing GC B cells was associated with impaired proliferation or survival. Unexpectedly, BrdU incorporation was not detectably reduced in GFP^+^ DGKζΔNLS-expressing GC B cells, indicating that enhanced DGKζ activity did not substantially impair their proliferation in vivo (Figure 5D). In contrast, DGKζΔNLS-expressing GC B cells showed increased cell death (Figure 5E). These findings indicate that the selective depletion of DGKζΔNLS-expressing GC B cells is associated with impaired survival rather than reduced proliferation.

We next determined whether enhanced DGKζ activity during the GC response compromised antigen-specific antibody production. Following immunization with the T-dependent antigen NP-CGG, EGFP-DGKζΔNLS-AID^Cre/+^ mice displayed increased total and high-affinity NP-specific IgM antibodies at day 7 after immunization. This increase was transient, with IgM titers becoming comparable to those of control mice at days 14 and 21. In contrast, total and high-affinity NP-specific IgG titers showed relatively modest reductions at day 7 but were markedly decreased at days 14 and 21 after immunization (Figure 5F). Thus, enhanced DGKζ activity during the GC response severely impaired the sustained NP-specific IgG response while largely preserving the IgM response.

Together, these results demonstrate that enhanced DGKζ activity selectively compromises the survival and accumulation of DGKζΔNLS-expressing GC B cells and severely impairs T-dependent antigen-specific IgG responses. Despite the selective depletion of DGKζΔNLS-expressing GC B cells, their proliferation was not detectably impaired in vivo, whereas cell death was increased, indicating that enhanced DGKζ activity primarily compromises GC B cell survival. These findings suggest that appropriately regulated DGK activity and DAG signaling are critical for maintaining GC B cells and generating robust antigen-specific IgG responses.

### DGKα/ζ deficiency promotes GC B cell expansion associated with enhanced signaling and glucose uptake

Our findings with DGKζΔNLS demonstrated that enhanced DGKζ activity and attenuation of DAG signaling impaired GC B cell accumulation and survival. We next asked whether perturbing DGK-mediated DAG metabolism in the opposite direction would alter GC B cell responses. Because mammalian cells express ten DGK isoforms (49), we first examined DGK family expression across B cell developmental and activation states using publicly available ImmGen Skyline RNA-seq data. DGK isoforms displayed distinct expression patterns across B cell development, with *Dgka, Dgkd*, and *Dgkz* among the more highly expressed isoforms in multiple B cell populations, including splenic GC centroblasts (CBs) and centrocytes (CCs) (Figure 6A and Supplemental Figure S2). Because previous studies suggested functional redundancy between DGKα and DGKζ in lymphocytes (28, 33, 35), we examined DGKα^-/-^ DGKζ^f/f^AID^Cre/+^ mice (hereafter DGKα/ζ DKO) to determine the consequences of combined DGKα and DGKζ deficiency during GC B cell responses.

**Figure 6.**
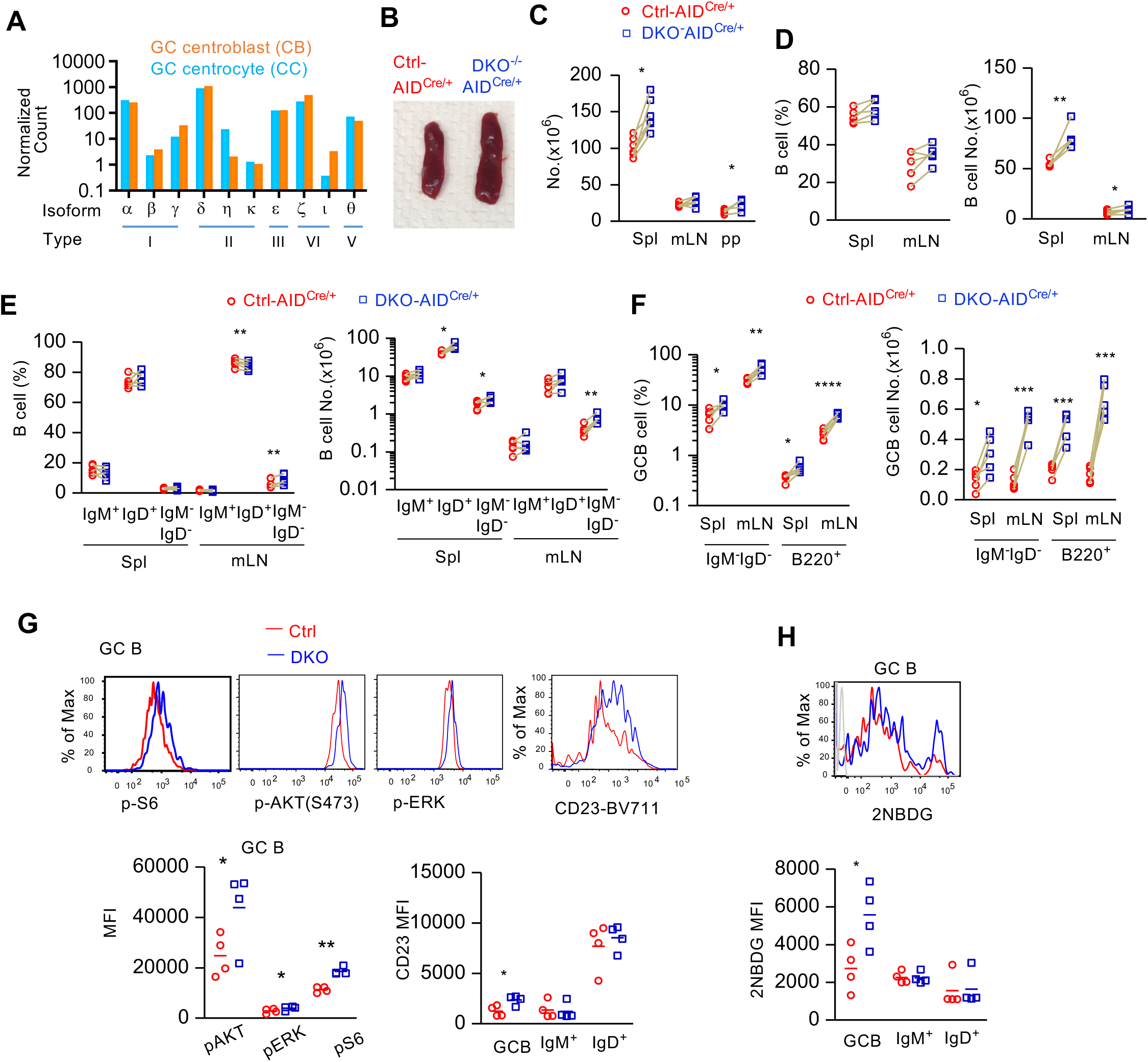
DGKα/ζ deficiency promotes GC B cell expansion associated with enhanced signaling and glucose uptake. (A) Normalized *Dgka, Dgkd,* and *Dgkz* mRNA expression in splenic germinal center (GC) centroblast (CB) and centrocyte (CC) populations from the ImmGen Skyline RNA-seq database. (B–H) Spleen, PP, and mLN cells from DGKα^-/-^ DGKζ^f/f^AID^Cre/+^ (DKO) and control mice were analyzed. (B) Representative spleens from control and DKO mice. (C) Total cell numbers in the spleen, mLN, and PPs. (D) Frequencies and absolute numbers of B220^+^ B cells in the spleen and mLNs. (E) Frequencies and absolute numbers of the indicated IgM/IgD-defined B cell populations in the spleen and mLNs. (F) Frequencies and absolute numbers of GC B cells analyzed within the B220^+^IgM^-^IgD^-^ population and within total B220^+^ B cells in the spleen and mLNs. (G) Representative histogram overlays and quantification of phosphorylated ERK1/2 (p-ERK1/2), Akt (p-Akt), and S6 (p-S6), together with CD23 expression, in GC B cells and the indicated non-GC B cell populations. (H) Representative histogram overlays and quantification of glucose uptake, measured using the fluorescent glucose analog 2-NBDG, in GC B cells and the indicated non-GC B cell populations. Each symbol represents one mouse. Data are presented as mean ± SEM where indicated and are representative of or pooled from three independent experiments. n.s., not significant; P < 0.05; P < 0.01; P < 0.001; P < 0.0001, as determined by unpaired two-tailed Student t-test.

DGKα/ζ DKO mice displayed visibly enlarged spleens compared with control mice (Figure 6B). Consistent with this observation, total cell numbers were increased in secondary lymphoid organs, including the spleen and PPs (Figure 6C). Although the frequencies of B cells were not substantially altered, their absolute numbers were increased in the spleen and PPs (Figure 6D). Analysis of B cell subsets revealed a prominent increase in IgM^-^IgD^-^ B cells, particularly in the spleen and mesenteric lymph nodes (mLNs) (Figure 6E). Most notably, GC B cells were markedly increased in both frequency and number in DGKα/ζ DKO mice (Figure 6F). These findings demonstrate that combined DGKα/ζ deficiency promotes a pronounced expansion of the GC B cell compartment.

We next examined whether the expanded GC B cell population was associated with alterations in intracellular signaling and cellular metabolism. DGKα/ζ-deficient GC B cells displayed enhanced phosphorylation of ERK1/2, Akt, and S6 compared with control cells, together with increased expression of CD23 (Figure 6G). Because Akt–mTOR signaling is closely linked to the regulation of cellular metabolism, including glucose uptake and utilization, we assessed glucose uptake using the fluorescent glucose analog 2-NBDG. DGKα/ζ-deficient GC B cells showed substantially increased 2-NBDG uptake compared with control cells (Figure 6H). In contrast, 2-NBDG uptake by IgM^+^ and IgD^+^ non-GC B cell populations was not consistently altered. Thus, the expansion of DGKα/ζ-deficient GC B cells was associated with enhanced signaling and increased glucose uptake.

Together, these findings reveal a phenotype opposite to that observed following enhanced DGKζ activity. Whereas enhanced DGKζ activity impaired GC B cell accumulation, combined DGKα/ζ deficiency promoted marked expansion of GC B cells in secondary lymphoid organs. The expanded DGKα/ζ-deficient GC B cell population exhibited enhanced ERK1/2, Akt, and S6 signaling together with increased glucose uptake, supporting a role for DGKα and DGKζ in restraining the signaling and metabolic state of activated GC B cells.

### DGKα/ζ deficiency promotes spontaneous autoreactivity but impairs antigen-specific antibody responses

The marked expansion of GC B cells in DGKα^-/-^DGKζ^f/f^AID^Cre^ mice prompted us to examine whether these changes were accompanied by altered antibody production. At steady state, serum IgM was increased in DGKα/ζ-deficient mice, whereas total IgG and the IgG subclasses examined were not broadly elevated (Figure 7A). Thus, GC B-cell accumulation did not translate into a generalized increase in serum IgG.

**Figure 7.**
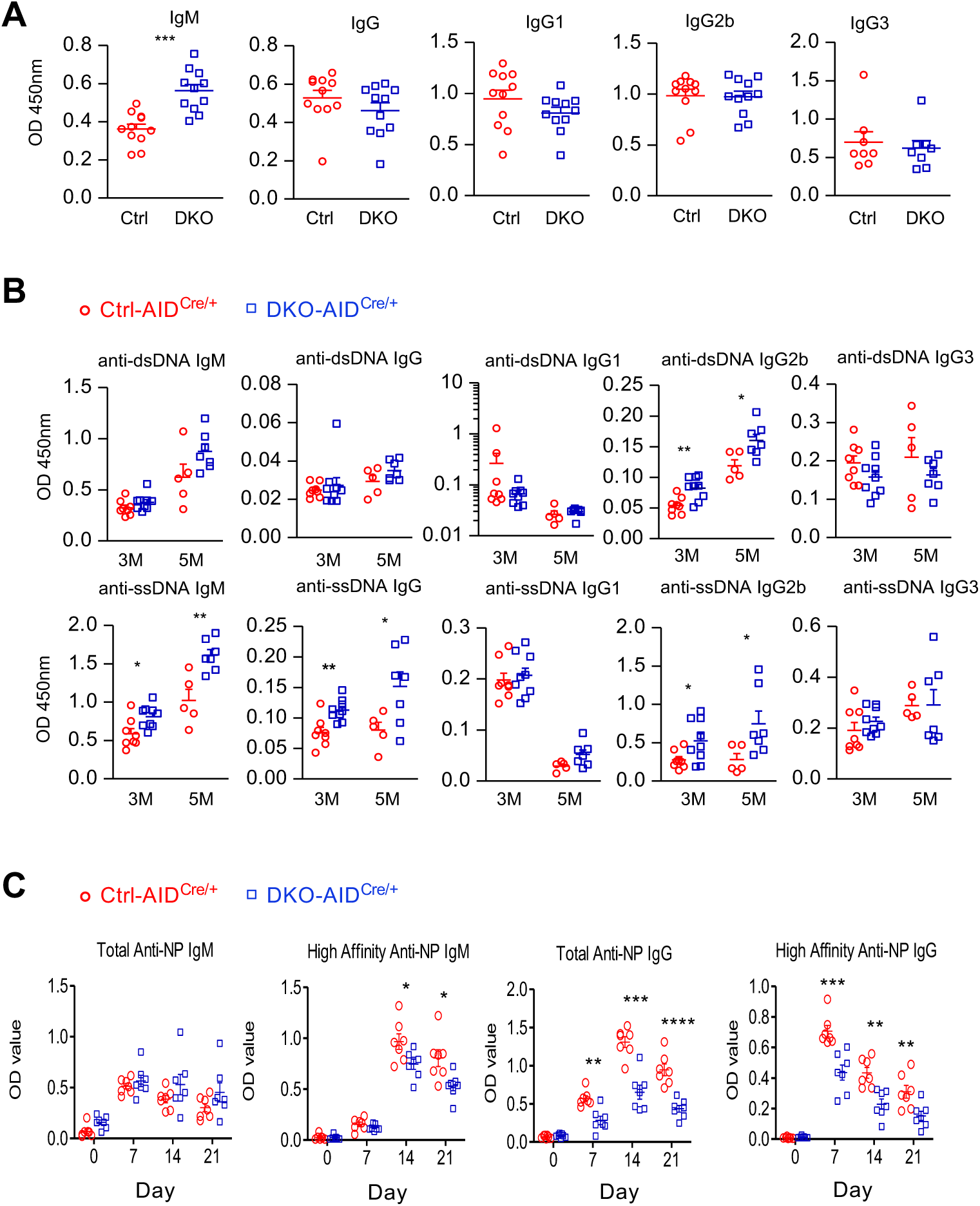
DGKα/ζ deficiency promotes spontaneous autoreactivity but impairs antigen-specific antibody responses. Serum antibodies from control and DGKα^-/-^DGKζ^f/f^-AID^Cre/+^ mice were measured by ELISA. (A) Steady-state serum IgM, total IgG, and IgG subclasses. (B) Anti-dsDNA and anti-ssDNA antibodies of the indicated isotypes at the indicated ages. (C) NP-specific total and high-affinity IgM and IgG responses following NP-CGG immunization, measured using NP26-BSA and NP4-BSA, respectively. Each symbol represents one mouse. Data are mean ± SEM pooled from two independent experiments. n.s., not significant; *P < 0.05; **P < 0.01; ***P < 0.001; ****P < 0.0001, determined by unpaired two-tailed Student t-test.

Because dysregulated GC responses can promote autoreactivity, we next measured anti-DNA antibodies. Anti-dsDNA antibodies remained relatively low and were not consistently increased, whereas anti-ssDNA antibodies were elevated in DGKα/ζ-deficient mice, particularly with age and across multiple isotypes (Figure 7B). These findings indicate that DGKα/ζ deficiency promotes selective spontaneous autoreactivity rather than a broad anti-DNA response.

We next assessed antigen-specific humoral responses following immunization with the T-dependent antigen NP-CGG. Despite expanded GC B-cell and Tfh compartments at steady state, DGKα/ζ-deficient mice showed impaired NP-specific antibody responses. Total NP-specific IgM was relatively preserved, whereas high-affinity IgM was reduced (Figure 7C). More strikingly, both total and high-affinity NP-specific IgG responses were substantially diminished compared with controls.

Together, these findings reveal a dissociation between GC B-cell accumulation and productive humoral immunity. DGKα/ζ deficiency promotes spontaneous ssDNA autoreactivity while impairing antigen-specific IgG responses, indicating that appropriate regulation of DAG signaling is required not only to restrain GC B-cell accumulation but also to support productive antigen-driven humoral responses.

## DISCUSSION

In this study, we demonstrate that DGK-mediated regulation of lipid signaling exerts stage-dependent effects on B cell development and GC responses. Enhanced DGKζ activity caused relatively modest perturbations during early BM B cell development but progressively impaired peripheral maturation, with pronounced effects on transitional and mature B cell populations. DGKζΔNLS also attenuated BCR-induced ERK1/2 and mTOR pathway activation and inhibited B cell activation and proliferation. Within the GC compartment, DGKζΔNLS selectively impaired GC B cell accumulation, primarily through reduced survival rather than a detectable defect in proliferation, and compromised antigen-specific IgG responses. Conversely, combined DGKα/ζ deficiency resulted in marked spontaneous GC B cell expansion associated with increased ERK1/2 and mTOR activation and enhanced glucose uptake. These mice nevertheless exhibited impaired antigen-specific antibody responses. Together, these findings suggest that DGKs establish an appropriate range of signaling required for B cell maturation and productive humoral responses, with both excessive attenuation and insufficient restraint of DGK-regulated signaling being detrimental.

The differential effects of DGKζΔNLS across developmental stages suggest that sensitivity to DAG signaling changes during B cell maturation. Pre-BCR and BCR engagement activates PLCγ2, which hydrolyzes PIP2 to generate IP3 and DAG, thereby coupling antigen-receptor signaling to Ca2+- and DAG-dependent pathways (24, 50). Signaling through the pre-BCR is required for the pro-B to pre-B transition (6, 7), and combined deletion of Erk1 and Erk2 causes a profound block at this transition by impairing pre-BCR-driven expansion and associated transcriptional programs (51). In comparison, DGKζΔNLS caused only a modest competitive disadvantage at the pro-B to pre-B transition and little apparent impairment of subsequent early BM developmental transitions. This milder phenotype may reflect incomplete attenuation of DAG signaling by DGKζΔNLS or contributions of DAG-independent pathways to ERK1/2 activation downstream of the pre-BCR. Consistent with the latter possibility, combined RasGRP1 and RasGRP3 deficiency markedly impairs BCR-induced Ras–ERK1/2 signaling and proliferation while having relatively modest effects on conventional B cell development (21). Thus, DGK-regulated DAG signaling contributes to the pre-BCR checkpoint but is unlikely to account for all signaling required for this developmental transition.

The consequences of enhanced DGKζ activity became considerably more pronounced during peripheral B cell maturation. In mixed BM chimeras, the competitive disadvantage of DGKζΔNLS-expressing cells increased significantly at the T1-to-T2 transition and became particularly pronounced among mature FO and MZ B cells, demonstrating a substantial cell-intrinsic component to this phenotype. Both canonical and noncanonical NF-κB pathways contribute to peripheral B cell maturation and maintenance, while BCR-generated DAG can activate PKCβ, which couples BCR signaling to the CARMA1–BCL10–MALT1 complex and canonical NF-κB activation (52, 53). Together with Ras–ERK1/2 signaling, these DAG-responsive pathways may contribute to the strong peripheral phenotype, although their individual contributions were not directly examined here. These findings indicate that peripheral B cell maturation is substantially more sensitive to altered DGK activity than most early BM developmental transitions.

Our signaling studies further indicate that DGKζ regulates both ERK1/2 and mTOR pathways in B cells. In T cells, DAG promotes mTOR activation through RasGRP1–Ras– ERK1/2- and PKCθ–CARMA1-dependent pathways, and DGKα and DGKζ restrain this response (31, 47). Similarly, enhanced DGKζ activity in B cells attenuated BCR-induced phosphorylation of both ERK1/2 and S6K1 and strongly inhibited BCR-induced CD69 expression and proliferation. Interestingly, DGKζΔNLS also impaired LPS-induced proliferation, particularly at lower LPS concentrations, whereas LPS-induced CD69 expression was largely preserved. Thus, enhanced DGKζ activity can influence proliferative responses beyond those initiated by the BCR while having less effect on some early TLR4-induced activation events. Because RasGRP1/RasGRP3 deficiency impairs BCR-induced proliferation while largely preserving LPS-driven proliferation (21), the effect of DGKζΔNLS on LPS responses may involve DGK-regulated pathways beyond RasGRP–Ras–ERK1/2signaling.

The GC compartment was particularly sensitive to enhanced DGKζ activity. Expression of DGKζΔNLS in AID-expressing B cells resulted in selective depletion of DGKζΔNLS-expressing cells from the GC compartment, whereas cells expressing kinase-inactive DGKζ were readily represented, demonstrating that this disadvantage depended on DGK catalytic activity. BrdU incorporation was not detectably reduced in DGKζΔNLS GC B cells, whereas cell death was increased, suggesting that enhanced DGKζ activity primarily compromises GC B cell survival. Consistent with impaired GC function, DGKζΔNLS mice exhibited markedly reduced NP-specific IgG responses despite relatively preserved IgM responses. ERK1/2 signaling also contributes to productive GC output, as GC B cell-specific ablation of Erk1/2 impairs antigen-specific plasma cell generation and antibody production, in part through defective induction of the plasma cell differentiation program (54). Thus, attenuation of Ras–ERK1/2 signaling may contribute to the impaired humoral response caused by DGKζΔNLS, although the accompanying reduction in S6K1 phosphorylation argues against attributing the phenotype exclusively to ERK1/2.

The preservation of GC B cell proliferation in vivo contrasts with the pronounced inhibition of BCR-driven proliferation by DGKζΔNLS in vitro. This difference may reflect the distinct signaling environment of GC B cells. BCR signaling is substantially rewired in GC B cells, where antigen-receptor signals are integrated with CD40, cytokine, and other microenvironmental inputs (14, 40, 41). These additional signals may sustain cell-cycle progression despite enhanced attenuation of BCR-DAG signaling while leaving survival more sensitive to altered DGK-DAG activity. Alternatively, changes in proliferation restricted to particular phases of the GC response may not have been detected by the BrdU approach used here. Nevertheless, our results identify impaired survival, rather than a generalized inability to proliferate, as the major detectable defect of DGKζΔNLS-expressing GC B cells.

Combined DGKα/ζ deficiency produced a striking reciprocal GC phenotype. Multiple DGK isoforms are expressed in B cells, with Dgka, Dgkd, and Dgkz among the more highly expressed isoforms in GC centroblasts and centrocytes. Nevertheless, combined DGKα/ζ deficiency caused marked spontaneous GC B cell expansion, consistent with overlapping functions of DGKα and DGKζ in restraining lymphocyte signaling (28, 31, 33–35). The relatively high expression of *Dgkd* suggests that DGK isoforms are not fully interchangeable, potentially reflecting differences in localization, regulation, or access to distinct DAG pools. DGKα/ζ-deficient GC B cells displayed increased ERK1/2 and mTOR activation together with increased 2-NBDG uptake, providing a biochemical counterpart to the attenuated ERK1/2 and S6K1 activation caused by enhanced DGKζ activity and linking DGK activity to the signaling and metabolic state of GC B cells.

mTORC1 is required for GC B cell proliferation and maintenance, and disruption of mTORC1 signaling impairs GC responses(55). However, increased mTORC1 activity alone is unlikely to explain the pronounced GC expansion caused by DGKα/ζ deficiency. Loss of TSC1 increases mTORC1 activity and disrupts B cell maturation but does not produce a comparable expansion of the GC compartment (56, 57). Together with increased ERK1/2 and Akt phosphorylation in DGKα/ζ-deficient GC B cells, these observations suggest that combined DGK deficiency alters a broader signaling network rather than acting solely through mTORC1. Similarly, increased 2-NBDG uptake indicates increased glucose uptake but does not establish increased glycolytic flux. How these signaling and metabolic changes contribute to the accumulation of DGKα/ζ-deficient GC B cells remains to be determined.

An important feature of the DGKα/ζ-deficient phenotype was the dissociation between spontaneous GC B cell expansion and productive antigen-specific humoral immunity. At steady state, DGKα/ζ-deficient mice exhibited markedly increased GC B cells accompanied by enhanced signaling and glucose uptake. In contrast, following NP-CGG immunization, these mice exhibited reduced high-affinity NP-specific antibody responses and markedly impaired NP-specific IgG production. Although GC responses were not directly examined following immunization, these findings indicate that spontaneous GC expansion does not predict an enhanced response to defined antigen. Dysregulated DGK-dependent signaling could affect affinity-dependent selection, class-switch recombination, or differentiation into antibody-secreting and memory B cells, although these possibilities require further investigation. The accompanying increase in serum IgM and age-associated anti-ssDNA antibodies further suggests that loss of DGK-mediated restraint favors spontaneous autoreactivity while compromising productive antigen-specific responses.

Several considerations limit attribution of these phenotypes exclusively to altered DAG abundance. DGKs both terminate DAG signaling and generate PA, which itself functions as a lipid second messenger; altered PA production could therefore contribute to the effects of either DGKζΔNLS expression or DGKα/ζ deficiency. In addition, the gain- and loss-of-function models are not exact genetic reciprocals: DGKζΔNLS enhances DGKζ activity, whereas the DKO combines germline DGKα deficiency with AID-Cre-dependent deletion of DGKζ. Nevertheless, the opposing effects of these manipulations on GC B cell accumulation, together with reciprocal changes in ERK1/2 and mTOR-associated signaling, strongly support an important role for DGKs in establishing the signaling state of GC B cells.

Collectively, our findings support a model in which DGKs tune lipid second-messenger signaling across B cell differentiation and in which sensitivity to this regulation varies with developmental stage. Early BM development is comparatively resistant to enhanced DGKζ activity, whereas peripheral maturation and GC B cell homeostasis are substantially more sensitive. Within the GC, excessive DGK activity compromises B cell survival and antigen-specific IgG responses, whereas insufficient DGK-mediated restraint promotes spontaneous GC B cell accumulation and heightened signaling and glucose uptake while also compromising productive antigen-specific antibody responses. Thus, productive humoral immunity requires DGK-regulated signaling to be maintained within an appropriate range rather than simply minimized or maximized.

## Acknowledgements

We thank the Flow Cytometry Core Facility of Duke Cancer Institute for the service and ChatGPT (OpenAI) for assistance with language editing and improving the clarity of the manuscript.

## Funding

Research in this manuscript is supported by the NIAID (R56AI079088, R01AI143781, R21AI200130, and R03AI195794).

## Conflict Of Interest

The authors declare no existing conflict of interest.

## Author Contributions

JX: Investigation, Methodology, Formal analysis, Visualization. SZ: Investigation, Formal analysis, Visualization. WL: Writing – review & editing. HT: Investigation. YP: Investigation. XL: Writing – review & editing. XPZ: Conceptualization, Methodology, Formal analysis, Supervision, Writing – original draft, Writing – review & editing.

## METHODS

### Mice

The conditional DGKζΔNLS and DGKξKD mouse strains were generated by targeting an EGFP-DGKζΔNLS or EGFP-DGKζKD expression cassette to the Rosa26 locus. The targeted allele contains a loxP-flanked transcriptional STOP cassette upstream of EGFP-DGKζΔNLS or EGFP-DGKζKD, allowing Cre-dependent expression of the fusion protein (45). Expression of the recombinant alleles were identified by GFP fluorescence. B6.Cg-Commd10Tg^(Vav1-icre)A2Kio^/J (Vav1iCre, stock no. 008610), B6.129P2(C)-Cd19^tm1(cre)Cgn^/J (CD19-Cre, stock no. 006785), and B6.129P2-Aicda^tm1(cre)Mn^z/J (AID-Cre, stock no. 007770) mice were obtained from The Jackson Laboratory. DGKα-deficient (Dgka^−/−^) mice and mice carrying a conditional DGKζ allele (Dgkz^f/f^) were reported previously (30, 33). Unless otherwise indicated, mice were analyzed at approximately 8–10 weeks of age. Experimental and control animals were age- and sex-matched, and both male and female mice were used. Mice were maintained under specific pathogen-free conditions. All animal procedures were performed in accordance with protocols approved by the Duke University Institutional Animal Care and Use Committee.

### Preparation of lymphoid cells

Single-cell suspensions were prepared from spleen, bone marrow (BM), Peyer’s patches (PPs), and mesenteric lymph nodes (mLNs), as indicated. BM cells were flushed from femurs and tibias with IMDM. Spleens, PPs, and mLNs were mechanically dissociated through 70-μm cell strainers. Erythrocytes in BM and spleen samples were lysed with ACK lysis buffer. Cells were washed and resuspended in IMDM containing 10% FBS and 50 μM 2-mercaptoethanol (2-ME). Viable cells were enumerated by trypan blue exclusion.

### Flow cytometry

Single-cell suspensions in 2% FBS-PBS were stained with the indicated fluorochrome-conjugated antibodies. Antibodies against B220, CD93, CD43, IgM, IgD, CD21, CD23, GL7, Fas, CD11b, Ly6C, Ly6G, Siglec-F, CD45.1, CD45.2, CD69, and BrdU were obtained from BioLegend, BD Biosciences, or Thermo Fisher Scientific. Antibody clones, fluorochromes, vendors, and catalog numbers are provided in Supplemental Table 1.

Unless otherwise indicated, cells were gated on singlets and viable cells. BM precursor and immature B cells were identified as B220^int^CD93^+^ and mature recirculating B cells as B220^hi^CD93^−^. Within the B220^int^CD93^+^ compartment, pro-B cells were identified as CD43^+^IgM^−^IgD^−^ and pre-B cells as CD43^−^IgM^−^IgD^−^. Large and small pre-B cells were distinguished by forward scatter (FSC^hi^ and FSC^lo^, respectively), and immature B cells were identified as CD93^+^IgM^+^IgD^−.^ Splenic transitional B cells were identified within the B220^int^CD93^+^ population. IgM^+^IgD^-^ T1 and T2 cells were distinguished by CD21/CD23 (T1: CD21^-^CD23^-^; T2: CD21^+^CD23^+^). Mature splenic B cells were identified within the B220^hi^CD93^−^ population. MZ B cells were defined as IgM^+^IgD^-^CD21^+^, and FO B cells as IgM^-^ IgD^+^CD23^+^CD21^lo^. GC B cells were identified as GL7^+^Fas^+^ cells within live B220^+^CD93^-^ IgM^−^IgD^−^ B cells. Monocytes (Ly6C^+^), neutrophils (Ly6G^+^), and eosinophils (SiglecF^+^) in mixed BM chimera experiments were identified using CD11b together with Ly6C, Ly6G, and Siglec-F. Cell viability was assessed using LIVE/DEAD Fixable Violet (Invitrogen) or 7-AAD as indicated. Samples were acquired on BD FACSCanto II or LSRFortessa flow cytometers and analyzed using FlowJo software (BD Biosciences; version 9).

### Mixed BM chimeras

CD45.1^+^ WT recipient mice were lethally irradiated with a single dose of 1,000 rad. Irradiated recipients were reconstituted by retro-orbital injection with a 1:1 mixture of BM cells from CD45.2^+^ WT and CD45.2^+^ DGKζΔNLS-Vav1^iCre^ donor mice. A total of 20 × 10^6^ BM cells was transferred per recipient. Eight weeks after reconstitution, BM and spleen cells were collected and analyzed by flow cytometry. Donor-derived cells were identified as CD45.1^−^CD45.2^+^, and WT and DGKζΔNLS-expressing donor populations were distinguished by GFP fluorescence. Competitive representation was calculated as the ratio of GFP^+^ DGKζΔNLS-expressing cells to GFP^−^ WT cells within each indicated B-cell or myeloid population.

### B-cell activation and proliferation

Splenocytes from DGKζΔNLS-CD19^Cre/+^ and WT-CD19^Cre/+^ control mice were labeled with CellTrace Violet (CTV; Thermo Fisher Scientific) according to the manufacturer’s instructions and cultured in complete IMDM containing 10% FBS, L-glutamine, penicillin/streptomycin, and 50 μM β-mercaptoethanol. Cells were left unstimulated or stimulated with the indicated concentrations of anti-IgM or LPS. For proliferation assays, cells were cultured for 80 h and proliferation was assessed by CTV dilution. Because CD19Cre-mediated recombination was incomplete, GFP^+^ DGKζΔNLS-expressing and GFP^−^ non-recombined B cells from the same cultures were analyzed separately and compared with control B cells. For analysis of early activation, splenocytes were stimulated overnight with anti-IgM or LPS at the indicated concentrations. Cells were then stained for B-cell markers and CD69, and CD69 expression was analyzed in GFP^+^, GFP^−^, and control B cells.

### BCR-induced signaling and immunoblotting

Splenic B cells from DGKζΔNLS-CD19^Cre/+^ and WT-CD19^Cre/+^ control mice were rested in PBS at 37°C for 30 minutes before stimulation. Cells were left unstimulated or stimulated with 10 μg/ml anti-IgM μ chain F(ab’)2 for 5 or 15 min at 37°C. Cells were lysed in 1% NP-40 lysis buffer containing 150 mM NaCl and 50 mM Tris-HCl (pH 7.4) supplemented with protease and phosphatase inhibitors. Cell lysates were resolved by SDS-PAGE and transferred to nitrocellulose membranes. Membranes were probed with antibodies against phospho-ERK1/2 (T200/Y204), total ERK1/2, phospho-S6K1 (T389), and β-actin. Bound antibodies were detected with HRP-conjugated secondary antibodies and enhanced chemiluminescence.

### Intracellular phospho-flow cytometry

Intracellular signaling was analyzed in freshly isolated cells from DGKα/ζ DKO and control mice. Cells were fixed with BD PhosFlow Fix Buffer I, permeabilized with BD Phosflow™ Perm/Wash Buffer I, and stained with antibodies specific for phospho-ERK1/2 (T200/Y204), phospho-Akt (S473), and phospho-S6 (S325/S326). GC B cells and the indicated IgM^+^ and IgD^+^ non-GC B-cell populations were identified by surface markers as described above. Phosphorylation was quantified as MFI within each population.

### Glucose uptake

Glucose uptake was assessed using the fluorescent glucose analog 2-(N-(7-Nitrobenz-2-oxa-1, 3-diazol-4-yl) Amino)-2-Deoxyglucose (2-NBDG). Freshly isolated splenocytes were incubated with 100 μM 2-NBDG for 30 minutes at 37°C in IMDM-10. 2-NBDG uptake was stopped by removing culture medium and washed with pre-chilled PBS twice. Cells were stained for surface markers before analysis with flow cytometry.

### In vivo BrdU incorporation and GC B-cell death

Mice were administered BrdU (1.5mg, MP Biomedicals) by intraperitoneal injection. Sixteen hours later, spleen and mLN cells were collected and stained for surface markers to identify GC B cells. Cells were subsequently fixed, permeabilized, and stained for incorporated BrdU using BD Pharmingen™ BrdU Flow Kit according to the manufacturer’s instructions. BrdU incorporation was quantified among GFP^+^ GC B cells. Cell death was assessed in GFP^+^ GC B cells using 7-AAD. The frequency of dead cells was determined by flow cytometry.

### Immunization and measurement of NP-specific antibodies

Mice were immunized by a single intraperitoneal injection of 20 μg NP18-CGG adsorbed to alum as previously described (56). Blood was collected before immunization and on days 7, 14, and 21 after immunization. Serum was isolated and stored at −80°C until analysis. Serum NP-specific antibodies were measured by ELISA. Ninety-six-well plates were coated overnight at 4°C with NP-BSA containing high NP26 or low NP4 hapten densities to measure total and high-affinity NP-specific antibodies, respectively. Diluted serum samples were added to coated plates. Bound antibodies were detected using HRP-conjugated antibodies specific for mouse IgM or IgG (Southern Biotech). Reactions were developed with TMB (Biolegend), and absorbance was measured at OD 450nm.

### Measurement of steady-state serum immunoglobulins

Steady-state serum IgM, total IgG, and the indicated IgG subclasses were measured by ELISA as previously described (56). Plates were coated with anti-mouse Igκ and Igλ antibodies (2 μg/ml; Southern Biotech) and incubated overnight at 4°C. Serially diluted serum samples were added, and bound immunoglobulins were detected using HRP-conjugated antibodies specific for mouse IgM, total IgG, or the indicated IgG subclasses. Absorbance was measured at OD 450 nm.

### Detection of anti-DNA autoantibodies

Serum anti-ssDNA and anti-dsDNA antibodies were measured by ELISA using sera collected from control and DGKα/ζ DKO mice at the indicated ages as previously described(35). Plates were coated with ssDNA or dsDNA and diluted serum samples were added. Bound antibodies were detected using HRP-conjugated antibodies specific for IgM, IgG, and indicated IgG isotypes. Reactions were developed with a TMB solution (Biolegend), and absorbance was measured at OD 450nm.

### Statistical analysis

Data are presented as mean ± SEM unless otherwise indicated. Comparisons between two independent groups were performed using two-tailed unpaired Student’s t-tests, whereas paired samples were analyzed using two-tailed paired Student’s t-tests. Statistical analyses were performed using GraphPad Prism version 10. P < 0.05 was considered statistically significant. Statistical significance is indicated as n.s., not significant; *P < 0.05; **P < 0.01; ***P < 0.001; and ****P < 0.0001.

## Notes

### Competing Interest Statement

The authors have declared no competing interest.

## REFERENCES

1. LeBien TW, Tedder TF. B lymphocytes: how they develop and function. Blood. 2008;112(5):1570–80.

2. Hardy RR, Hayakawa K. B cell development pathways. Annu Rev Immunol. 2001;19:595–621.

3. Melchers F. Checkpoints that control B cell development. J Clin Invest. 2015;125(6):2203–10.

4. Winkler TH, Mårtensson I-L. The Role of the Pre-B Cell Receptor in B Cell Development, Repertoire Selection, and Tolerance. Frontiers in Immunology. 2018;9:2423.

5. Zehentmeier S, Pereira JP. Cell circuits and niches controlling B cell development. Immunol Rev. 2019;289(1):142–57.

6. Gu H, Kitamura D, Rajewsky K. B cell development regulated by gene rearrangement: arrest of maturation by membrane-bound D mu protein and selection of DH element reading frames. Cell. 1991;65(1):47–54.

7. Kitamura D, Roes J, Kuhn R, Rajewsky K. A B cell-deficient mouse by targeted disruption of the membrane exon of the immunoglobulin mu chain gene. Nature. 1991;350(6317):423–6.

8. Kitamura D, Kudo A, Schaal S, Muller W, Melchers F, Rajewsky K. A critical role of lambda 5 protein in B cell development. Cell. 1992;69(5):823–31.

9. Loder F, Mutschler B, Ray RJ, Paige CJ, Sideras P, Torres R, et al. B cell development in the spleen takes place in discrete steps and is determined by the quality of B cell receptor-derived signals. J Exp Med. 1999;190(1):75–89.

10. Allman D, Srivastava B, Lindsley RC. Alternative routes to maturity: branch points and pathways for generating follicular and marginal zone B cells. Immunological Reviews. 2004;197:147–60.

11. Pillai S, Cariappa A. The follicular versus marginal zone B lymphocyte cell fate decision. Nature Reviews Immunology. 2009;9(11):767–77.

12. Berek C, Berger A, Apel M. Maturation of the immune response in germinal centers. Cell. 1991;67(6):1121–9.

13. Jacob J, Kelsoe G, Rajewsky K, Weiss U. Intraclonal generation of antibody mutants in germinal centres. Nature. 1991;354(6352):389–92.

14. Victora GD, Nussenzweig MC. Germinal Centers. Annual Review of Immunology. 2022;40:413–42.

15. Young C, Brink R. The unique biology of germinal center B cells. Immunity. 2021;54(8):1652–64.

16. Kurosaki T, Shinohara H, Baba Y. B cell signaling and fate decision. Annu Rev Immunol. 2010;28:21–55.

17. Packard TA, Cambier JC. B lymphocyte antigen receptor signaling: initiation, amplification, and regulation. F1000Prime Rep. 2013;5:40.

18. Ishiai M, Kurosaki M, Pappu R, Okawa K, Ronko I, Fu C, et al. BLNK required for coupling Syk to PLC gamma 2 and Rac1-JNK in B cells. Immunity. 1999;10(1):117–25.

19. Bell SE, Vigorito E, McAdam S, Reynolds HM, Caraux A, Colucci F, et al. PLCgamma2 regulates Bcl-2 levels and is required for survival rather than differentiation of marginal zone and follicular B cells. Eur J Immunol. 2004;34(8):2237–47.

20. Yu M, Chen Y, Zeng H, Zheng Y, Fu G, Zhu W, et al. PLCgamma-dependent mTOR signalling controls IL-7-mediated early B cell development. Nat Commun. 2017;8(1):1457.

21. Coughlin JJ, Stang SL, Dower NA, Stone JC. RasGRP1 and RasGRP3 regulate B cell proliferation by facilitating B cell receptor-Ras signaling. Journal of Immunology. 2005;175(11):7179–84.

22. Leitges M, Schmedt C, Guinamard R, Davoust J, Schaal S, Stabel S, et al. Immunodeficiency in protein kinase cbeta-deficient mice. Science. 1996;273(5276):788–91.

23. Wheeler ML, Dong MB, Brink R, Zhong XP, DeFranco AL. Diacylglycerol kinase zeta limits B cell antigen receptor-dependent activation of ERK signaling to inhibit early antibody responses. Sci Signal. 2013;6(297):ra91.

24. Wang D, Feng J, Wen R, Marine JC, Sangster MY, Parganas E, et al. Phospholipase Cgamma2 is essential in the functions of B cell and several Fc receptors. Immunity. 2000;13(1):25–35.

25. Hashimoto A, Takeda K, Inaba M, Sekimata M, Kaisho T, Ikehara S, et al. Cutting edge: essential role of phospholipase C-gamma 2 in B cell development and function. J Immunol. 2000;165(4):1738–42.

26. Wen R, Chen Y, Schuman J, Fu G, Yang S, Zhang W, et al. An important role of phospholipase Cgamma1 in pre-B-cell development and allelic exclusion. EMBO J. 2004;23(20):4007–17.

27. Zhong XP, Guo R, Zhou H, Liu C, Wan CK. Diacylglycerol kinases in immune cell function and self-tolerance. Immunol Rev. 2008;224:249–64.

28. Guo R, Wan CK, Carpenter JH, Mousallem T, Boustany RM, Kuan CT, et al. Synergistic control of T cell development and tumor suppression by diacylglycerol kinase alpha and zeta. Proc Natl Acad Sci U S A. 2008;105(33):11909–14.

29. Zhong XP, Hainey EA, Olenchock BA, Jordan MS, Maltzman JS, Nichols KE, et al. Enhanced T cell responses due to diacylglycerol kinase zeta deficiency. Nat Immunol. 2003;4(9):882–90.

30. Olenchock BA, Guo R, Carpenter JH, Jordan M, Topham MK, Koretzky GA, et al. Disruption of diacylglycerol metabolism impairs the induction of T cell anergy. Nat Immunol. 2006;7(11):1174–81.

31. Gorentla BK, Wan CK, Zhong XP. Negative regulation of mTOR activation by diacylglycerol kinases. Blood. 2011;117(15):4022–31.

32. Shin J, O’Brien TF, Grayson JM, Zhong XP. Differential regulation of primary and memory CD8 T cell immune responses by diacylglycerol kinases. J Immunol. 2012;188(5):2111–7.

33. Yang J, Zhang P, Krishna S, Wang J, Lin X, Huang H, et al. Unexpected positive control of NFkappaB and miR-155 by DGKalpha and zeta ensures effector and memory CD8+ T cell differentiation. Oncotarget. 2016;7(23):33744–64.

34. Yang J, Wang HX, Xie J, Li L, Wang J, Wan ECK, et al. DGK alpha and zeta Activities Control TH1 and TH17 Cell Differentiation. Front Immunol. 2019;10:3048.

35. Li L, Huang H, Wang H, Pan Y, Tao H, Zhang S, et al. DGKalpha and zeta deficiency causes regulatory T-cell dysregulation, destabilization, and conversion to pathogenic T-follicular helper cells to trigger IgG1-predominant autoimmunity. Elife. 2026;14.

36. Victora GD, Schwickert TA, Fooksman DR, Kamphorst AO, Meyer-Hermann M, Dustin ML, et al. Germinal Center Dynamics Revealed by Multiphoton Microscopy with a Photoactivatable Fluorescent Reporter. Cell. 2010;143(4):592–605.

37. Shulman Z, Gitlin AD, Weinstein JS, Lainez B, Esplugues E, Flavell RA, et al. Dynamic signaling by T follicular helper cells during germinal center B cell selection. Science. 2014;345(6200):1058–62.

38. Gitlin AD, Shulman Z, Nussenzweig MC. Clonal selection in the germinal centre by regulated proliferation and hypermutation. Nature. 2014;509(7502):637–40.

39. Gitlin AD, Mayer CT, Oliveira TY, Shulman Z, Jones MJ, Koren A, et al. HUMORAL IMMUNITY. T cell help controls the speed of the cell cycle in germinal center B cells. Science. 2015;349(6248):643–6.

40. Khalil AM, Cambier JC, Shlomchik MJ. B cell receptor signal transduction in the GC is short-circuited by high phosphatase activity. Science. 2012;336(6085):1178–81.

41. Luo W, Weisel F, Shlomchik MJ. B Cell Receptor and CD40 Signaling Are Rewired for Synergistic Induction of the c-Myc Transcription Factor in Germinal Center B Cells. Immunity. 2018;48(2):313–26.e5.

42. Mueller J, Matloubian M, Zikherman J. Cutting edge: An in vivo reporter reveals active B cell receptor signaling in the germinal center. J Immunol. 2015;194(7):2993–7.

43. Gonzalez DG, Cote CM, Patel JR, Smith CB, Zhang Y, Nickerson KM, et al. Nonredundant Roles of IL-21 and IL-4 in the Phased Initiation of Germinal Center B Cells and Subsequent Self-Renewal Transitions. J Immunol. 2018;201(12):3569–79.

44. Tanaka T, Okada M, Hozumi Y, Tachibana K, Kitanaka C, Hamamoto Y, et al. Cytoplasmic localization of DGKzeta exerts a protective effect against p53-mediated cytotoxicity. J Cell Sci. 2013;126(Pt 13):2785–97.

45. Xie D, Zhang S, Chen P, Deng W, Pan Y, Xie J, et al. Negative control of diacylglycerol kinase zeta-mediated inhibition of T cell receptor signaling by nuclear sequestration in mice. Eur J Immunol. 2020;50(11):1729–45.

46. Rickert RC, Roes J, Rajewsky K. B lymphocyte-specific, Cre-mediated mutagenesis in mice. Nucleic Acids Res. 1997;25(6):1317–8.

47. Hamilton KS, Phong B, Corey C, Cheng J, Gorentla B, Zhong X, et al. T cell receptor-dependent activation of mTOR signaling in T cells is mediated by Carma1 and MALT1, but not Bcl10. Sci Signal. 2014;7(329):ra55.

48. Robbiani DF, Bothmer A, Callen E, Reina-San-Martin B, Dorsett Y, Difilippantonio S, et al. AID is required for the chromosomal breaks in c-myc that lead to c-myc/IgH translocations. Cell. 2008;135(6):1028–38.

49. Krishna S, Zhong XP. Regulation of Lipid Signaling by Diacylglycerol Kinases during T Cell Development and Function. Front Immunol. 2013;4:178.

50. Oh-hora M, Johmura S, Hashimoto A, Hikida M, Kurosaki T. Requirement for Ras guanine nucleotide releasing protein 3 in coupling phospholipase C-gamma2 to Ras in B cell receptor signaling. J Exp Med. 2003;198(12):1841–51.

51. Yasuda T, Sanjo H, Pages G, Kawano Y, Karasuyama H, Pouyssegur J, et al. Erk kinases link pre-B cell receptor signaling to transcriptional events required for early B cell expansion. Immunity. 2008;28(4):499–508.

52. Guldenpfennig C, Teixeiro E, Daniels M. NF-kB’s contribution to B cell fate decisions. Front Immunol. 2023;14:1214095.

53. Sommer K, Guo B, Pomerantz JL, Bandaranayake AD, Moreno-Garcia ME, Ovechkina YL, et al. Phosphorylation of the CARMA1 linker controls NF-kappaB activation. Immunity. 2005;23(6):561–74.

54. Yasuda T, Kometani K, Takahashi N, Imai Y, Aiba Y, Kurosaki T. ERKs induce expression of the transcriptional repressor Blimp-1 and subsequent plasma cell differentiation. Sci Signal. 2011;4(169):ra25.

55. Li B, Li Z, Wang P, Huang Q, Xu L, He R, et al. Mammalian target of rapamycin complex 1 signalling is essential for germinal centre reaction. Immunology. 2017;152(2):276– 86.

56. Ci X, Kuraoka M, Wang H, Carico Z, Hopper K, Shin J, et al. TSC1 Promotes B Cell Maturation but Is Dispensable for Germinal Center Formation. PLoS One. 2015;10(5):e0127527.

57. Benhamron S, Tirosh B. Direct activation of mTOR in B lymphocytes confers impairment in B-cell maturation andloss of marginal zone B cells. Eur J Immunol. 2011;41(8):2390–6.

