## Supplemental Figure S1 for "Diacylglycerol Kinases Balance DAG Signaling to Control B Cell Maturation, Germinal Center Homeostasis, and Humoral Immunity"

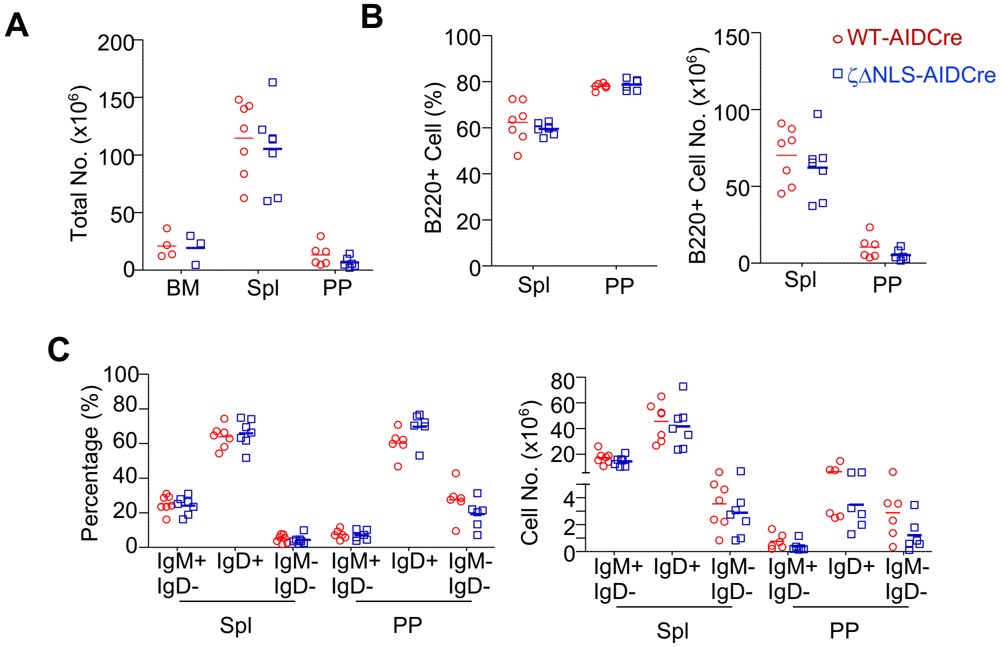


**Supplemental Figure S1. Analysis of B cell populations in DGKζΔNLS-AIDCre and DGKζKD-AIDCre mice.** BM, spleen, and Peyer's patch (PP) cells from DGKζΔNLS-AID$^{Cre/+}$ and control mice were analyzed. (A) Total cell numbers in the BM, spleen, and PPs. (B) Frequencies and absolute numbers of B220$^+$ B cells. (C) Frequencies and absolute numbers of IgM$^+$IgD$^-$, IgM$^-$IgD$^+$, and IgM$^-$IgD$^-$ B-cell populations. Data were pooled from at least three independent experiments.
