## Supplemental Figure S2 for "Diacylglycerol Kinases Balance DAG Signaling to Control B Cell Maturation, Germinal Center Homeostasis, and Humoral Immunity"

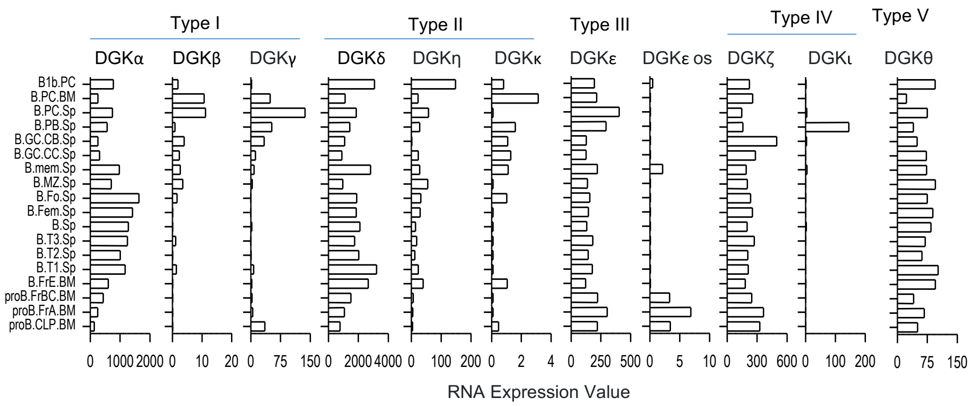


| Population Name | Population Long Name | Description |
|---|---|---|
| proB.CLP.BM | Bone Marrow Common Lymphoid Progenitor | CD93+CD117+IL7Ra+CD45R- |
| proB.FrA.BM | Bone Marrow Fr. A (Pre-Pro-B) | CD93+CD117+IL7Ra+CD45R+ |
| proB.FrBC.BM | Bone Marrow Fr. BC (Pro-B) | CD93+IgM-CD19+CD43+HSA+ |
| B.FrE.BM | Bone Marrow Fr. E (newly formed B) | CD93+IgM+CD19+CD43-HSA+ |
| B.T1.Sp | Splenic T1 | CD19+CD45R+IgM++CD93+CD23- |
| B.T2.Sp | Spleinc T2 | CD19+CD45R+IgM++CD93+CD23+ |
| B.T3.Sp | Splenic T3 | CD19+CD45R+IgM+CD93+CD23+ |
| B.Sp | Splenic B cells | CD19+Igm+TCRb- |
| B.Fem.Sp | Female Splenic B cells | CD19+Igm+TCRb- |
| B.Fo.Sp | Splenic Follicular | CD19+CD45R+IgM+CD93-CD23+CD43-CD5- |
| B.MZ.Sp | Splenic Marginal Zone | CD19+CD45R+IgM++CD93-CD23-CD21/35++ |
| B.mem.Sp | Splenic Memory | sorted on CD19+B220+IgD-Fas-CD38+IgG+ |
| B.GC.CC.Sp | Splenic Germinal Center Centrocytes | CD19+B220+IgD-Fas+CD38-CXCR4lo CD86hi |
| B.GC.CB.Sp | Splenic Germinal Center Centroblasts | CD19+B220+IgD-Fas+CD38-CXCR4hi CD86lo |
| B.PB.Sp | Splenic Plasmablasts | CD138+ MHCIIhi (dump: NK1.1- TCRb- Gr1- CD11b-) |
| B.PC.Sp | Splenic Plasma cells | BLIMP1+ CD138+ MHCIIlo (dump: NK1.1- TCRb- Gr1- CD11b-) |
| B.PC.BM | Bone Marrow Plasma cells | BLIMP1+ CD138+ MHCIIlo (dump: NK1.1- TCRb- Gr1- CD11b-) |
| B1b.PC | Peritoneal B1b | CD19+CD45R+IgM++CD93-CD23-CD43+CD5- |

6 week old C57BL/6J mice

**Supplemental Figure S2. Expression of DGK family members across B-cell developmental and activation states**. RNA expression of the indicated diacylglycerol kinase (DGK) isoforms across FACS-purified B-cell populations was obtained from the ImmGen Skyline RNA-seq dataset. B-cell populations are ordered according to developmental or activation state and include bone marrow progenitors (proB.CLP.BM, proB.FrA.BM, proB.FrBC.BM, and B.FrE.BM); splenic transitional B cells (B.T1.Sp, B.T2.Sp, and B.T3.Sp); mature splenic B-cell populations (B.Sp, B.Fem.Sp, B.Fo.Sp, and B.MZ.Sp); memory B cells (B.mem.Sp); germinal center centrocytes (B.GC.CC.Sp) and centroblasts (B.GC.CB.Sp); splenic plasmablasts and plasma cells (B.PB.Sp and B.PC.Sp); bone marrow plasma cells (B.PC.BM); and peritoneal B1b cells (B1b.PC). Expression values represent normalized RNA-seq values reported by ImmGen Skyline.
